# Classification of RNA backbone conformation into rotamers using ^13^C′ chemical shifts: How far we can go?

**DOI:** 10.1101/559302

**Authors:** A. A. Icazatti, J.M. Loyola, I. Szleifer, J.A. Vila, O. A. Martin

## Abstract

The conformational space of the ribose–phosphate backbone is very complex as is defined in terms of six torsional angles. To help delimit the RNA backbone conformational preferences 46 rotamers have been defined in terms of the these torsional angles. In the present work, we use the ribose experimental and theoretical ^13^C′ chemical shifts data and machine learning methods to classify RNA backbone conformations into rotamers and families of rotamers. We show to what extent the use of experimental ^13^C′ chemical shifts can be used to identify rotamers and discuss some problem with the theoretical computations of ^13^C′ chemical shifts.

## INTRODUCTION

Nucleic acids are central macromolecules for the storing, flow and regulation of genetic and epigenetic information in cellular organisms. RNA can adopt a wide variety of 3D structural conformations and this structural variability explains the multiplicity of roles that RNA performs on cells (1, 2). The classification of RNA backbone conformations into rotamers is a very useful way to delimit the conformational space of RNA structures. Rotamers are defined in terms of the backbone torsional angles namely *α*, *β*, *γ*, *δ*, *ε*, *ζ* (as shown in Figure 1). This classification was proposed by Richardson et al 2008 (3), and has been achieved after the attempts of different research groups to find a consensus RNA backbone structural classification. There are 55 backbone rotamers, from which 46 are rotamers with well defined torsional angles distributions, and the remaining 9 rotamers were proposed as *wannabe* rotamers. The ‘suite’ is the basic subunit used for rotamer classification. The suite is defined from sugar-to-sugar (or from the *δ* torsional angle of residue *i-1* to the *δ* torsional angle of residue *i*), and it is contained within the dinucleotide subunit (see Figure 1).

**Figure 1.**
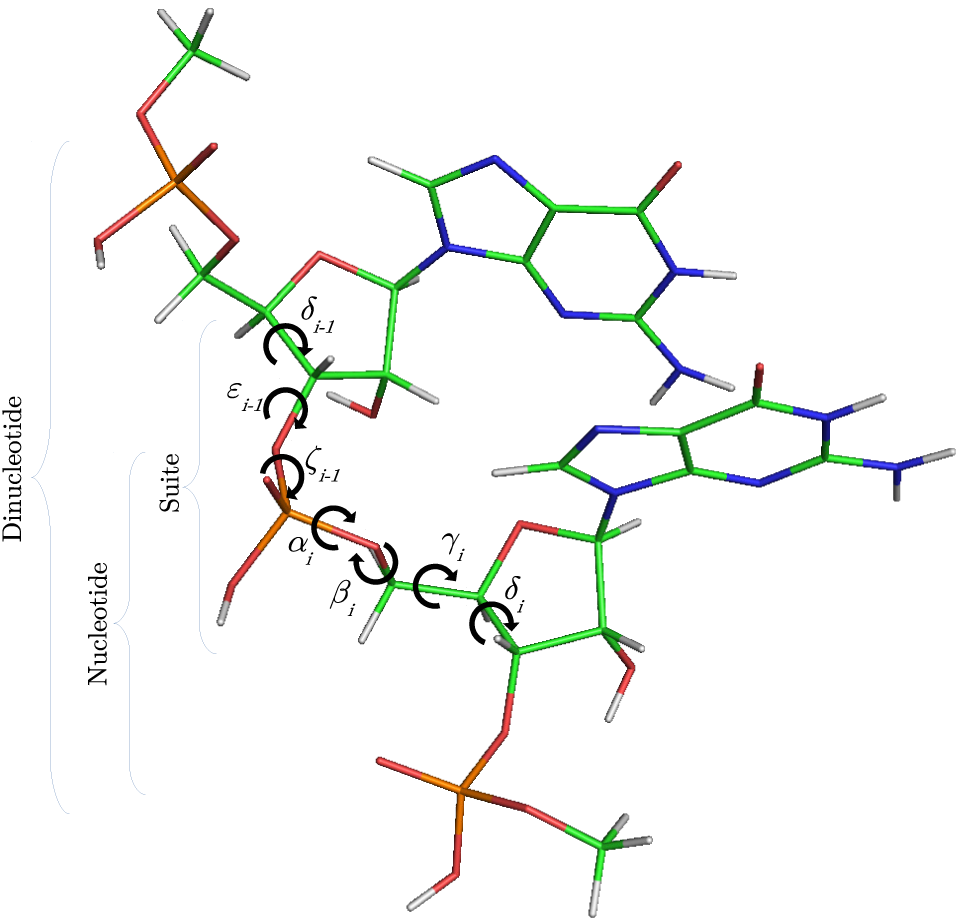
RNA dinucleotide. C, H, O, N and P nuclei are colored in green, white, red, blue and orange, respectively. Torsional angles of RNA backbone are named on Greek characters (*α*, *β*, *γ*, *δ*, *ε*, *ζ*). Suite (from *δ*_*i*−1_ to *δ*_*i*_), dinucleotide and nucleotide subunits are indicated.

^13^C′ chemical shifts have been successfully used by our and other groups for protein and glycan structural determination, validation and refinement (4, 5, 6, 7, 8). In this work, we study how to use ^13^C′ chemical shifts to classify RNA backbone into rotamers.

## MATERIALS AND METHODS

A dataset of RNA backbone rotamers with ^13^C′ chemical shifts values is necessary to train the machine learning models to classify RNA experimental suites into rotamers. In the following two section we explain how we obtained two datasets.

### Experimental dataset

Experimental ^13^C′ chemical shift data for RNA molecules was retrieved from the BioMagResBank (BMRB; www.bmrb.wisc.edu)(9), along with their corresponding structures from the Protein Data Bank (PDB; https://www.rcsb.org/) (10). As it is fundamental to count on reliable experimental ^13^C′ chemical shifts values for an accurate structural analysis, data curation was carried out using 13Check RNA (11) a python module to correct RNA ^13^C′ chemical shifts systematic errors, recently developed in our group. The obtained dataset (see Supplementary Table S1) contains 26 RNA structures with ^13^C′ chemical shifts for the five ribose carbon nuclei (C1′, C2′, C3′, C4′ and C5′), providing a total of 391 suite subunits. Given that we needed a one-to-one correspondence between the sets of chemical shifts and the rotamer suites, only the first structure from each NMR ensemble was used, considering that the first model listed in the PDB files is usually reported as the structure with the lowest energy scoring. For every PDB entry, the 3D coordinates of the first model were extracted in order to compute the backbone torsional angles (*δ*_*i* − 1_, *ε*_i − 1_, *ζ*_i − 1_, *α*_i_, *β_i_*, *γ_i_*, *δ_i_*) of the suites. Then, these torsional angles were used to assign the RNA suites to their corresponding rotamer names. From the 46 original rotamers, only 38 are represented in the final experimental dataset.

### Theoretical dataset

In order to have a complete dataset with the 46 RNA backbone rotamers and their corresponding ^13^C′ chemical shifts, a theoretical dataset was also constructed. A template for each of the 16 possible combinations of dinucleotide (A, C, U and G) sequences was obtained from RNA structures found in the PDB. A Monte-Carlo conformational sampling was carried out by rotating the backbone torsional angles of the corresponding suite contained in each dinucleotide, given the torsional angle distributions for each of the 46 RNA backbone rotamer suites (3) (the 9 *wannabe* rotamers were excluded from this analysis) while keeping the bond-lengths and bond-angles fixed (rigid geometry approximation). As a result, 10,340,852 conformations were generated. Quantum-theory level computation of chemical shifts is very time-consuming. Therefore, to reduce the number of calculations, a smaller number of conformations was selected. Aiming to keep most of the variability of the originally generated conformations, we computed the Shannon entropy (S) (see Equation 1) of the distribution of torsional angles. The entropy was computed for different subsets of conformations and sample sizes (from 5 to 100) (see Figure 2). We decided to use the 80% of the maximum entropy as a cutoff, which implies around 40 conformations per rotamer. As we also considered the 16 combinations of dinucleotide sequences, the total number of conformations computed at the DFT level of theory was 30,530.

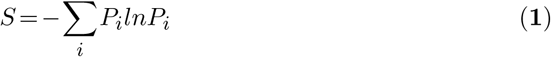

**Figure 2.**
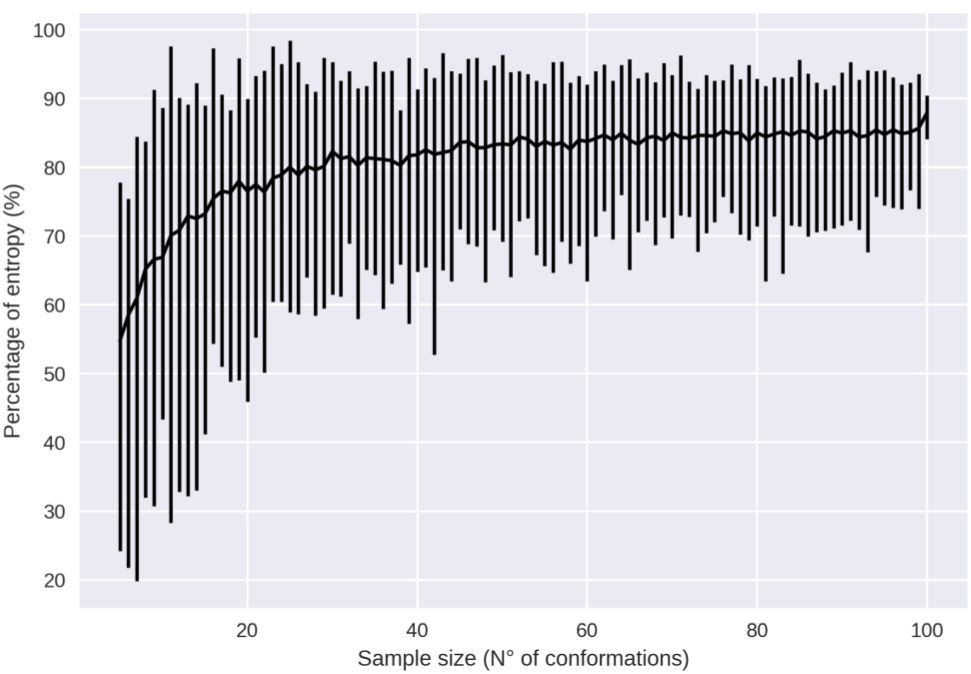
Percentage of entropy of the sample against sample size for a given dinucleotide sequence and rotamer, UU and 1a, respectively, in this case.

### Details of the quantum-chemical calculations of the ^13^C′ shieldings

To perform the DFT calculations, the obtained dinucleotide conformations were split in their corresponding mononucleotide subunits. A test showed that when mononucleotides were used instead of dinucleotides, the result was exactly the same within 10^−2^ppm and the total computation time was approximately half the total time for computing the complete dinucleotides. Nucleotide subunits were treated as terminally-blocked mononucleotides with methyl groups (Me) in both termini 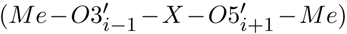. Phosphate groups of the backbone were treated as neutral, because we assume that all backbone charges are shielded during the quantum-chemical calculations. This approach was adopted because under physiological conditions, the phosphate groups are completely ionized and neutralized by positive charges (13). A 6–311+G(2d,p) locally dense basis set (14) was used for calculation of backbone ^13^C′ chemical shifts and their nearest neighbour nuclei, at the DFT level of approximation (see Figure 3 for details). The remaining nuclei were treated with a 3-21G basis set. The OB98 density functional was used because good results were previously observed for proteins and glycans in our group (15, 16). All DFT computations were done using the Gaussian package (12).

**Figure 3.**
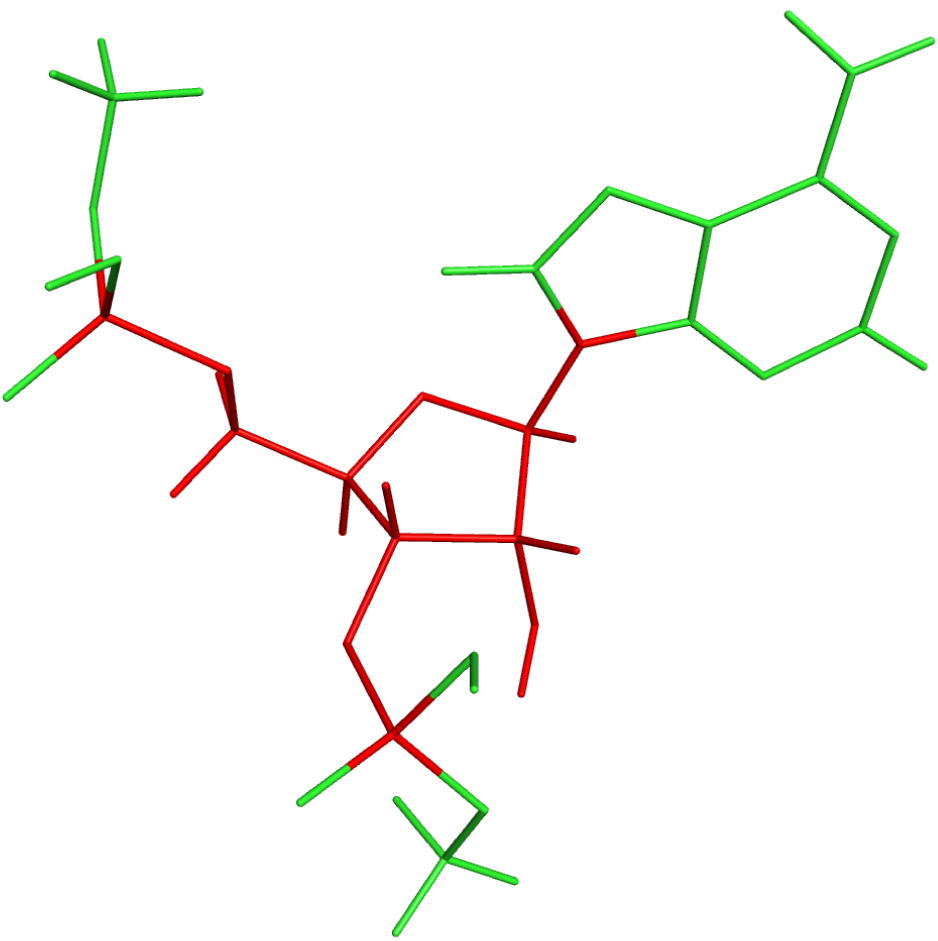
Example of a methyl blocked mononucleotide used for DFT calculations. The locally-dense basis-set approach is indicated by the different colors: the nuclei in red were treated with the extended 6-311+G(2d,p) basis set and the nuclei in green were treated with the smaller 3-21G basis set.

### Families of rotamers

The original 46 RNA backbone rotamers were grouped in families based on their *δ*_*i* − 1_, *δ_i_*, *α* and *γ* torsional angles values. Only these 4 (out of 7) backbone torsional angles in the suite subunit were chosen to group the rotamers because their distributions of observed values are bimodal (*δ*_*i* − 1_ and *δ_i_*) and trimodal (*γ* and *α*), with clearly separated peaks (see Figure 4), which allowed us to group rotamers based on the torsional angle values within the different peaks. As summarized in Table 1, 4 families were found when both *δ*_*i* − 1_ and *δ*_*i* −_ torsional angles in the suite were used, 7 families for the *αγ* combination, 10 families for *δ*_*i* − 1_*δ*_*i*_α, and *δ*_*i* − 1_*δ_i_γ*, and 22 families for *δ*_*i* − 1_*δ*_*i*_*αγ*. In order to evaluate the classification performance of the RNA A–form helix conformations, the rotamers were also grouped as: (i) A_noA families, where the 46 rotamers were separated in A–form helix (1a) vs. no A–form helix rotamers, and (ii) A*_noA* families, where the 46 rotamers were separated in rotamers related to A–form helix (1a, 3d, 3b, 5d, 0a, 6b and 4b rotamers) vs. the remaining rotamers.

**Table 1.** Families of rotamers

| 46<br>rotamers | 22 families<br>$\delta_{i-1}\delta_i\alpha\gamma$ | 10 families<br>$\delta_{i-1}\delta_i\alpha$ | 10 families<br>$\delta_{i-1}\delta_i\gamma$ | 7 families<br>$\alpha\gamma$ | 4 families<br>$\delta_{i-1}\delta_i$ | 2 families<br>A_noA<sup>i</sup> | 2 families<br>A*_noA<sup>*ii</sup> |
| --- | --- | --- | --- | --- | --- | --- | --- |
| &a | e | a | a | e | a | b | b |
| #a | q | c | c | e | c | b | b |
| 0a | q | c | c | e | c | b | a |
| 0b | t | d | d | e | d | b | b |
| 0i | o | g | g | b | c | b | b |
| 1[ | l | b | b | e | b | b | b |
| 1a | e | a | a | e | a | a | a |
| 1b | l | b | b | e | b | b | b |
| 1c | d | e | e | d | a | b | b |
| 1e | f | e | e | f | a | b | b |
| 1f | d | e | e | d | a | b | b |
| 1g | c | a | a | c | a | b | b |
| 1L | e | a | a | e | a | b | b |
| 1m | e | a | a | e | a | b | b |
| 1o | m | i | i | g | b | b | b |
| 1t | k | f | f | d | b | b | b |
| 1z | j | b | b | c | b | b | b |
| 2[ | t | d | d | e | d | b | b |
| 2a | q | c | c | e | c | b | b |
| 2h | r | g | g | f | c | b | b |
| 2o | v | j | j | g | d | b | b |
| 3a | e | a | a | e | a | b | b |
| 3b | l | b | b | e | b | b | a |
| 3d | a | a | a | a | a | b | a |
| 4a | q | c | c | e | c | b | b |
| 4b | t | d | d | e | d | b | a |
| 4d | n | c | c | a | c | b | b |
| 4g | p | c | c | c | c | b | b |
| 4n | o | g | g | b | c | b | b |
| 4p | s | d | d | a | d | b | b |
| 4s | u | h | h | f | d | b | b |
| 5d | a | a | a | a | a | b | a |
| 5j | b | e | e | b | a | b | b |
| 5q | h | f | f | b | b | b | b |
| 5z | j | b | b | c | b | b | b |
| 6d | n | c | c | a | c | b | a |
| 6g | p | c | c | c | c | b | b |
| 6j | o | g | g | b | c | b | b |
| 6n | o | g | g | b | c | b | b |
| 6p | s | d | d | a | d | b | b |
| 7a | e | a | a | e | a | b | b |
| 7d | a | a | a | a | a | b | b |
| 7p | g | b | b | a | b | b | b |
| 7r | i | i | i | c | b | b | b |
| 8d | n | c | c | a | c | b | b |
| 9a | e | a | a | e | a | b | b |

The 46 RNA backbone rotamers were arranged in 22, 10, 10, 7 and 4 families of rotamers based on the observed distributions of $\delta_{i-1}\delta_i\alpha\gamma$ , $\delta_{i-1}\delta_i\alpha$ , $\delta_{i-1}\delta_i\gamma$ , $\alpha\gamma$ and $\delta_{i-1}\delta_i$ torsional angles values, respectively. Additionally, the 46 rotamers were separated in RNA A-form helix vs. no A-form helix rotamers in two ways: (i) RNA A-form helix rotamer 1a vs. the remaining no A-form helix rotamers (A\_noA families) and (ii) rotamers related to A-form helix (i.e. 1a, 3d, 3b, 5d, 0a, 6b, 4b) vs. the remaining rotamers (A\*\_noA\* families).

**Figure 4.**
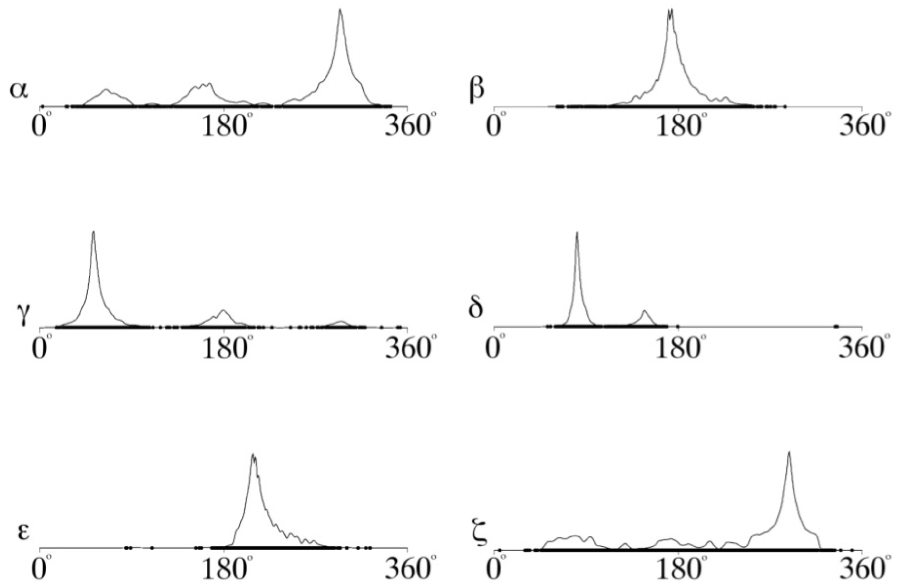
RNA backbone torsional angles distributions. Reproduced with authors permission from Laura Weston Murray (2007) “RNA Backbone Rotamers and Chiropraxis” Doctoral Dissertation; Dept. of Biochemistry; Duke University, 169 pages. Chaper 2, figure 6.

### Classification

A series of machine learning methods were used to classify RNA suites as rotamers (or families of rotamers) based on their ribose ^13^C′ chemical shifts values. The following classification methods from the scikit-learn Python library (17) were trained: K-Nearest Neighbors (NN), Decision Tree (DT), Random Forest (RF), Support Vector Machine (SVM) and a class of neural network called Multi-Layer Perceptron (MLP). Different model parameters were tried out (see Supplementary Table S3). A random sampling algorithm was also used as a control, where suites were classified randomly. The sequence of the suite was considered for classification, because we found that the performance increased compared to a sequence–independent classification (see Supplementary Figure S1).

The classification performance was assessed with two measures: weighted accuracy and *F*_1_ score (21). The weighted accuracy was used in order to recalibrate the contribution of the different rotamers, because the observed frequency of the rotamers is highly uneven (e.g. the A–form helix rotamer 1a has an observed frequency of ~0.75).The weights used in the weighted accuracy were obtained from a substitution matrix (ROSUM, for ROtamers SUbstitution Matrix). The definition of the ROSUM matrix was inspired by the BLOSUM matrix used for protein sequence alignment (18). The matrix is used to weight the match or no match, between the true rotamer and the predicted rotamer, as a function of the euclidean distance between rotamers (in the seven-dimensional space of the suite backbone torsional angles) and the observed frequency of each rotamer. The torsional angles values and the observed frequencies are extracted from the rotamers table (3). A ROSUM matrix was obtained for each of the rotamer families described in the previous section (see Supplementary Data Section 2). The f1–score was also used as a performance measure because it is the harmonic mean of precision and recall (https://en.wikipedia.org/wiki/F1 score3) and as such, it gives “a more realistic measure of the classifier’s performance” (https://www.quora.com/Whats-the-advantage-of-using-the-F1-score-when-evaluating-classification-performance).

*Experimental vs theoretical* The classification models trained with theoretical data were used to classify the experimental suites. The result of the theoretical calculations (described in a previous section) are theoretical NMR isotropic shieldings (*σ*). The theoretical shieldings (*σ_comp_*) must be subtracted from a reference shielding value (*σ_ref_*) to be transformed into theoretical chemical shifts (*δ_comp_*) (see Equation 2) which can then be compared with the experimental chemical shifts (*δ_exp_*). A simple reference value of *σ_ref_*=185.00 ppm was used, which is very close to the theoretical isotropic shielding for TMS (*σ_T M S,th_*) (15), and it is consistent with the reference value previously defined for proteins and glycans.

Alternatively, a set of effective references were obtained as a function of: (i) the nitrogenous base sequence, (ii) the combinations of ribose puckering states in the four families of rotamers obtained from *δ*_*i*−1_*δ_i_* torsional angles distributions, (iii) the five carbon nuclei ^13^C′ CS mean values and (iv) a linear regression between theoretical and experimental ribose ^13^C′ CS values for a set of suites (see Supplementary Table S2).

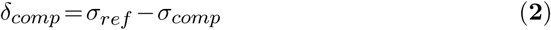

*Theoretical vs theoretical* The classification models trained with theoretical data were also used to classify the theoretical suites. In this case, classification was assessed through a leave-one-out cross-validation (LOO-CV). In LOO-CV, the dataset is split into a test set and training set in a one-folded manner, which means that at every iteration a unique suite is taken apart from the dataset and the remaining suites are used for training. This process continues until every suite from the theoretical dataset is evaluated.

*Experimental vs experimental* A LOO-CV was also used to classify the experimental suites.

## RESULTS AND DISCUSSION

For experimental vs theoretical classification (Figure 5a), the 46 rotamers can be classified by means of backbone ^13^C′ chemical shifts with a maximal *F*_1_ score of 0.34 (see Supplementary Table S5). When the 46 rotamers are grouped in families based on their torsional angles distributions, the highest scores correspond to the use of *δ*_(*i−*1)_and *δ*_(*i*)_ torsional angles, where all the classifiers gave maximal scores above 0.65. This result is in agreement with the fact that backbone ^13^C′ chemical shifts are highly sensitive to ribose puckering states (19), since the *δ* torsional angle keeps a direct relation with the ribose puckering (20). The *δ*_*i* −1_*δ_i_γ*, *δ*_*i* − 1_*δ*_*i*_α, *δ*_*i* −1_*δ_i_αγ* and *αγ* families also show improved scores over the classification of the 46 rotamers. The A* noA* and A_noA families show low classification scores relative to their random choice classification score, which means that backbone ^13^C′ chemical shifts cannot distinguish between A–form helix and no A–form helix rotamers. In general the use of more complex classifier models such as Neural Networks, Support Vector Machine, Decision Tree and Random Forest does not assure a better performance for the current task, thus the simpler Nearest Neighbor model can be chosen for classification into RNA rotamers. In both the theoretical dataset LOO-CV and the experimental dataset LOO-CV (see Figure 5b and 5c, respectively), the performance increase for every group of families, compared to the experimental vs theoretical classification. In the theoretical dataset LOO-CV the performance values are very close to 1.0 for *δ*_*i* −1_*δ_i_* families and A–form helix/no A–form helix rotamers (A_noA). In the theoretical dataset LOO-CV, the performance value ranges are particularly narrow, except for MLP and SVM classifiers.

**Figure 5.**
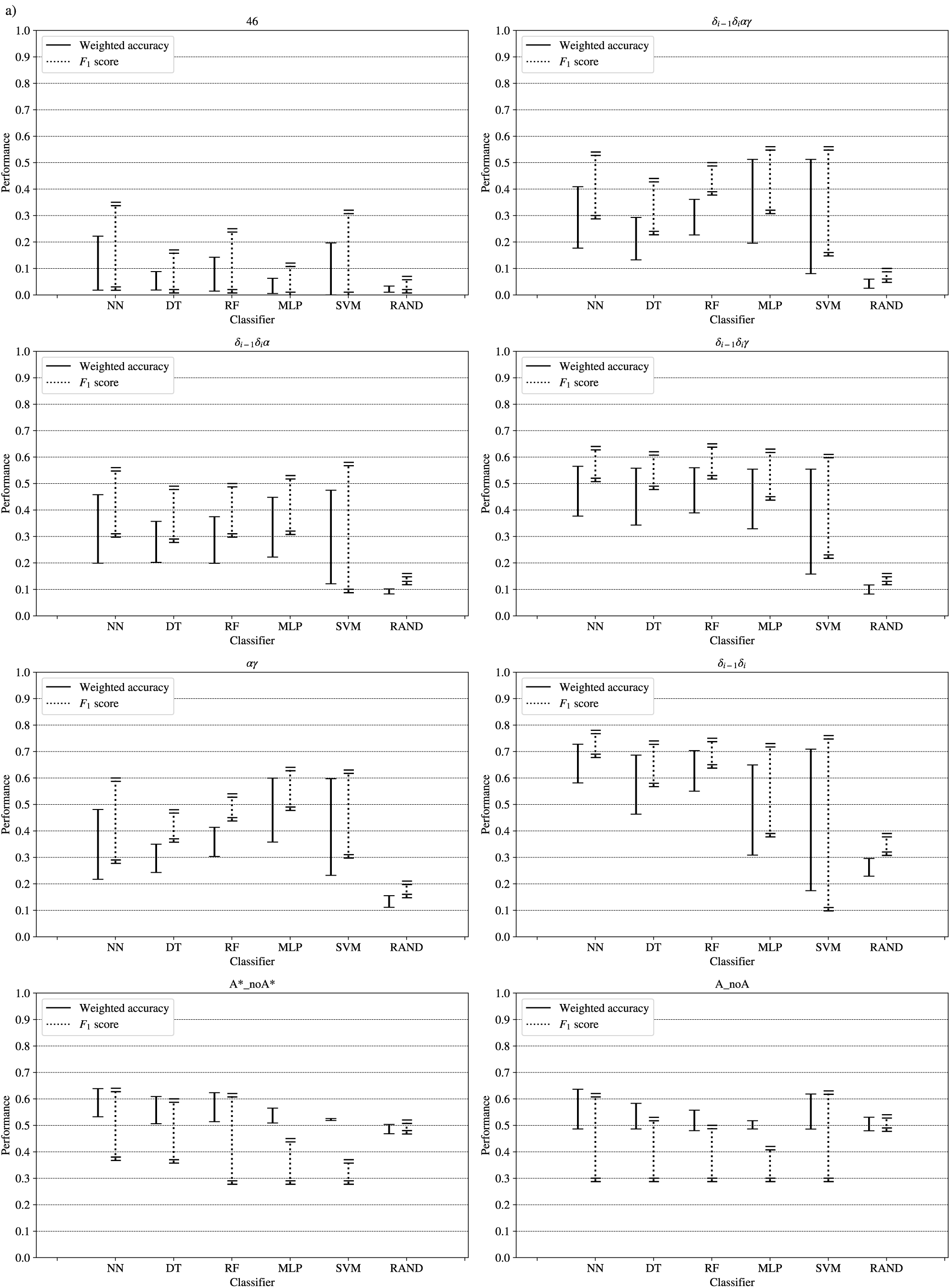
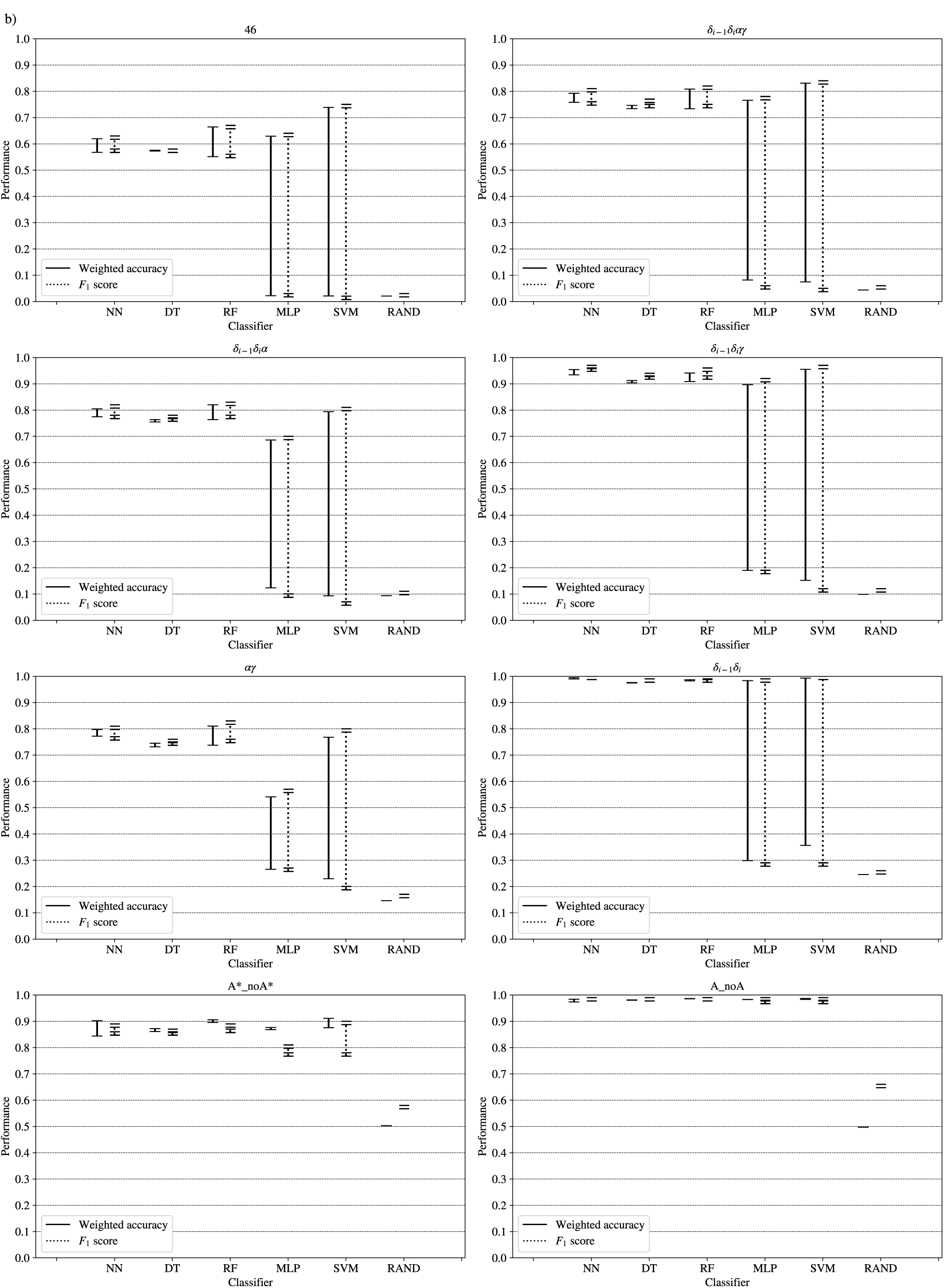
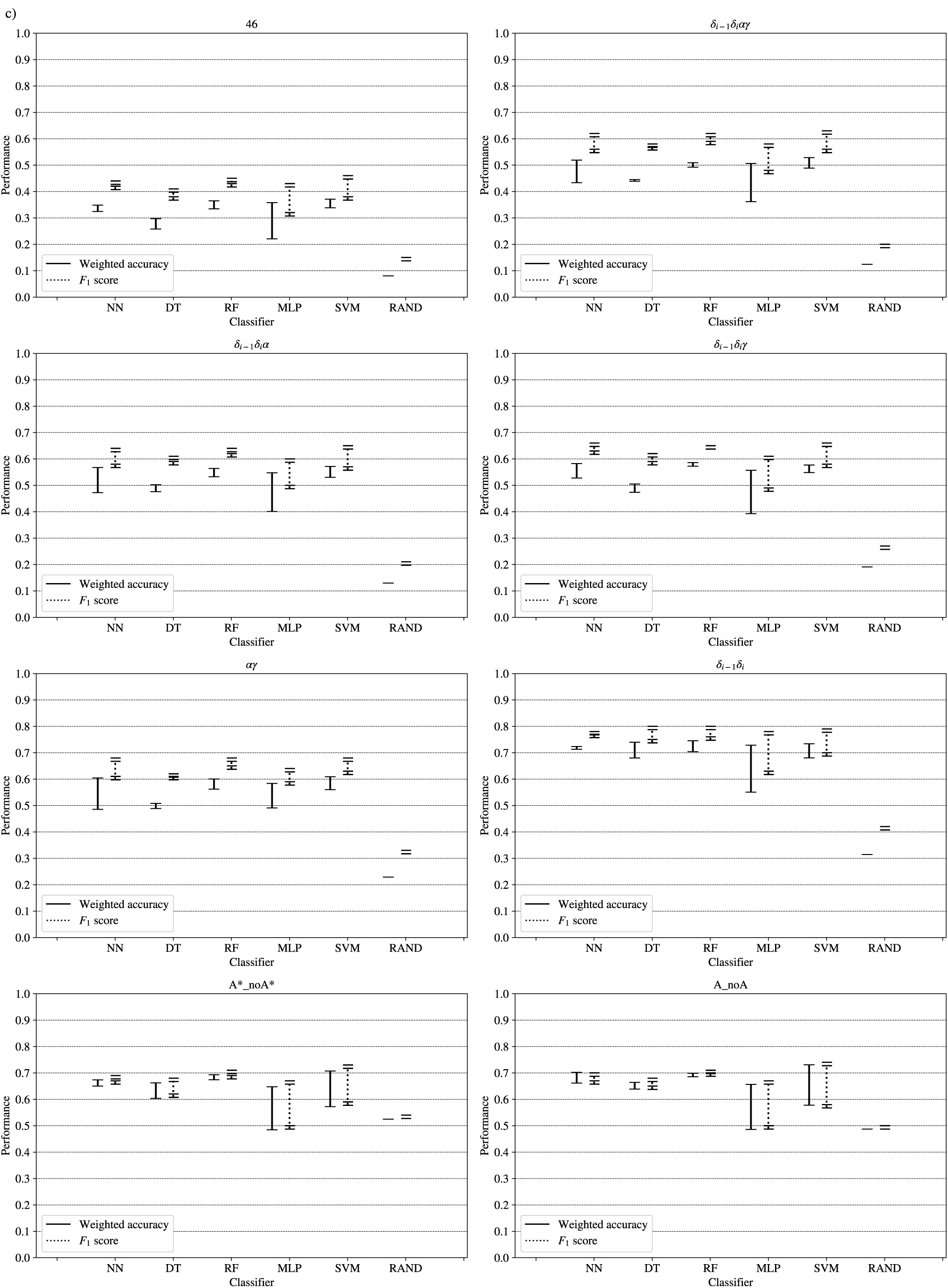
Value ranges of weighted accuracy and *F*_1_ score for the classification of rotamers and families of rotamers, using Nearest Neighbour (NN), Decision Tree (DT), Random Forest (RF), Multi-Layer Perceptron (MLP) and Support Vector Machine (SVM) classifiers. A random-choice (RAND) algorithm was used as a baseline reference. In a), the classification models were generated from theoretical data and were used to classify the experimental data. The results from theoretical dataset LOO-CV and experimental dataset LOO-CV are shown in b) and c), respectively. The highest values of weighted accuracy and *F*_1_ score, for the classification results shown in a), along with parameters of the classifiers are provided in Supplementary Tables S4 and S5.

The high scores obtained for the theoretical vs theoretical classification indicates that ^13^C′ chemical shifts are in fact very sensitive to changes of the torsional angles, the only variable we changed for the construction of the theoretical dataset. At the same time the lower performance obtained in the experimental vs theoretical classification, is signalling that the atomistic model used for the DFT computations is not good enough to reproduce the experimental observations.

One reason the theoretical vs theoretical classification gives better results compared to the experimental vs experimental classification could be that the experimental database is very sparse and the theoretical dataset is instead dense, or in other words the coverage of the theoretical dataset is much more better than the experimental one. To explore if this is in fact a reasonable explanation we remove elements from the theoretical dataset to mimic the sparsity of the experimental dataset. We found that while the accuracy decreased (on average 0.09 points) this is not enough to explain the lower performance of the experimental vs theoretical or experimental vs experimental classification. Reinforcing the idea discussed in the previous paragraph, i.e we need a better model for the theoretical DFT computations. This experiment also provides indirect evidence indicating that the accuracy of the experimental vs experimental classification will be improved as more RNA conformations are deposited in databases giving another incentive to determine and deposit RNA structures and ^13^C′ chemical shifts data.

## CONCLUSION

We explored, the use of RNA backbone ^13^C′ chemical shifts to classify backbone conformations into rotamers and families of rotamers. In general our study led us to the following conclusions: (1) The classification of the rotamer families defined by the *δ* torsional angles, which are directly related to the ribose puckering states, gives the best performance; in line with result previously described by other authors; (2) Classification of A-form helix and no A-form helix rotamers using ^13^C′ chemical shifts is not better than a random classification; (3) The accuracy achieved using the simple nearest-neighbour method is on par with more complex classifiers such as Neural Networks, Support Vector Machine, Decision Tree and Random Forest; (4) ^13^C′ chemical shifts values are able to sense change in torsional angles, but they are also affected by other factors, thus future DFT computations of RNA ^13^C′ chemical shifts should use more complex models than the one used in this work; (5) Experimental ^13^C′ chemical shifts can be useful to identify RNA rotamers, if the rotamers are re-grouped in smaller families as the 46 rotamers seems to be too fine description for accurate discrimination in terms of ^13^C′ chemical shifts; (6) the usefulness of ^13^C′ chemical shifts for rotamers identification should improve as more RNA structures and experimental ^13^C′ chemical shifts becomes available.

## ACKNOWLEDGEMENTS

We greatly appreciate Myriam Villegas for valuable discussions, comments and suggestions.

## FUNDING

This research was supported by grants from: Consejo Nacional de Investigaciones Científicas y Técnicas (Argentina) [PIP-0087 (JAV)], Agencia Nacional de Promoción Científica y Tecnológica (Argentina) [PICT-0556 and PICT-0767 (JAV) and PICT-0218 (OAM)].

## Conflict of interest statement

None declared.

