## Supplementary Data for "Classification of RNA backbone conformation into rotamers using ^13^C′ chemical shifts: How far we can go?"

#### 1 Experimental database

| PDB ID | BMRB ID |
|---|---|
| 1AJU | 15869 |
| 1HWQ | 5007 |
| 1KKA | 5256 |
| 1NC0 | 5655 |
| 1Q75 | 5932 |
| 1R7W | 6076 |
| 1YSV | 6485 |
| 1ZC5 | 6633 |
| 2JPP | 15257 |
| 2KOC | 5705 |
| 2KZL | 17316 |
| 2LU0 | 18503 |
| 2M22 | 18892 |
| 2M4W | 19024 |
| 2MEQ | 18975 |
| 2MFC | 19544 |
| 2MFF | 19547 |
| 2MFG | 19548 |
| 2MHI | 19634 |
| 2N7X | 25826 |
| 2N82 | 25831 |
| 2O32 | 15080 |
| 2QH2 | 7403 |
| 2QH3 | 7404 |
| 2QH4 | 7405 |
| 5KQE | 30132 |

Table S1: BMRB and PDB IDs for the 26 RNA structures of the experimental dataset

#### 2 Use of nucleotide sequence for classification

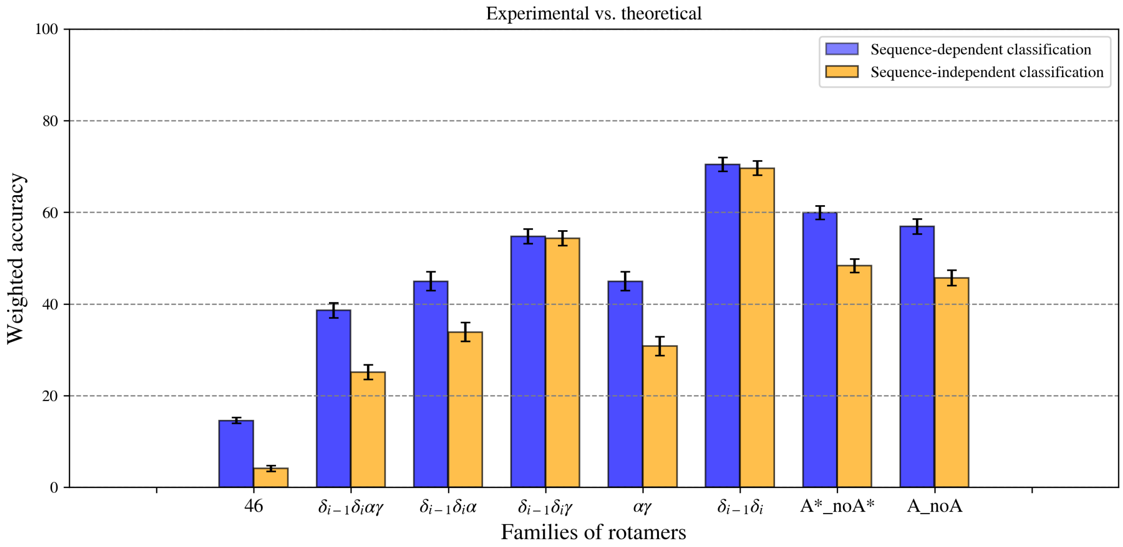


Supplementary Figure S1: Weighted accuracies for sequence-dependent and sequence-independent Nearest–Neighbor classifications, for experimental dataset against theoretical dataset. Groups labels A_noA, A*_noA* and 46 indicate A–form helix rotamer (1a) vs. no A–form helix rotamers, A–form helix related rotamers (1a, 3d, 3b, 5d, 0a, 6b and 4b) vs. no A–form helix related rotamers, and the 46 original rotamers. While families labels $\delta_{i-1}\delta_i$, $\alpha\gamma$, $\delta_{i-1}\delta_i\alpha$, $\delta_{i-1}\delta_i\gamma$, $\delta_{i-1}\delta_i\alpha\gamma$ indicate the combination of torsional angles used for grouping. Blue and orange bars indicate sequence–dependent and sequence–independent classifications, respectively. Error bars indicate the standard deviation in the weighted accuracies for the different conformations in the NMR assemblies.

#### 3 Construction of the ROSUM matrices

The $jk$-element of the ROSUM matrix (denoted as $a_{jk}$) is defined by Equation 1, where $P_{jk}$ is the probability of substitution of rotamer$_j$ by rotamer$_k$, and it is defined by Equation 2. The probability of substituion $P_{jk}$ is obtained from the normalized distance $D_{norm}$ (Equations 3 and 4), where distance $D_{jk}$ is the sum of the intra–rotameric distances $d_{jj}$ and $d_{kk}$, and the inter–rotameric distance $d_{jk}$. An intra–rotameric distance is the average of the torsion angle standard deviations of a given rotamer (Equations 5 and 6). The inter–rotameric distance is the euclidean distance between the torsion angles mean values of rotamer$_j$ and rotamer$_k$ (Equation 7). Both the torsion angles mean values and the standard deviations were extracted from the Richardson's rotamer table [1].

$$a_{jk} = log(\frac{P_{jk}}{q_j * q_k}) \quad (1)$$

$$P_{jk} = \frac{1}{D_{norm}} \quad (2)$$

$$D_{norm,s} = \frac{D_{jk}}{\sum D_{jk}} \quad (3)$$

$$D_{jk} = d_{jj} + d_{kk} + d_{jk} \quad (4)$$

$$d_{jj} = \frac{\sqrt{\sum (\sigma_{x_j})^2}}{7} \quad (5)$$

$$d_{kk} = \frac{\sqrt{\sum (\sigma_{x_k})^2}}{7} \quad (6)$$

$$d_{jk} = \sqrt{\sum (x_j - x_k)^2} \quad (7)$$

where:

$$x \quad = \delta_{i-1},\ \varepsilon_{i-1},\ \zeta_{i-1},\ \alpha_i,\ \beta_i,\ \gamma_i,\ \delta_i$$

When j and k are rotamer families, the frequencies $q_j$ and $q_k$ are the sum over the observed frequencies of the rotamer members of families j and k, respectively, and $D_{norm}$ is the average distance between the rotamer members of families j and k.

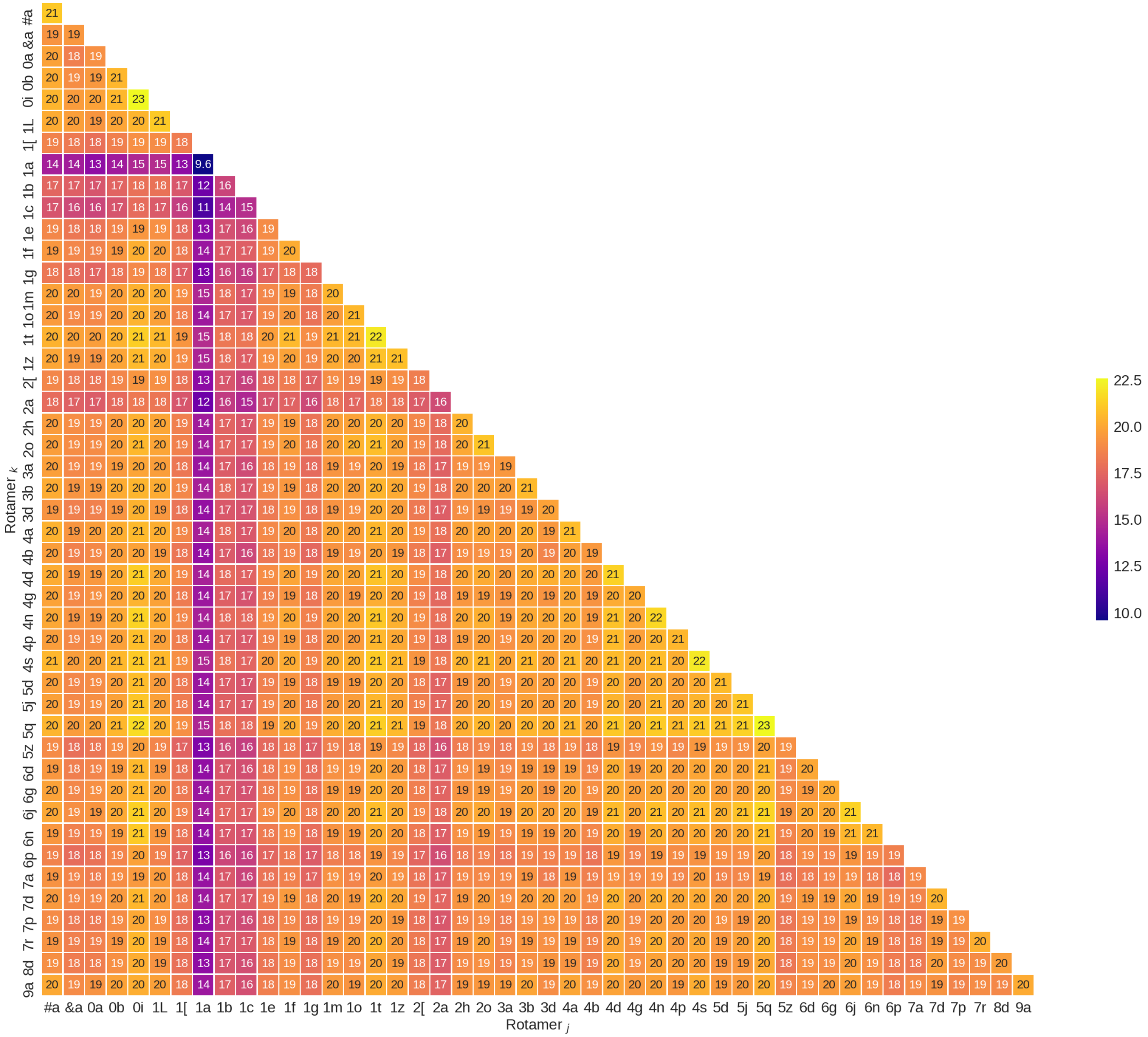


Figure S2: Representation of the ROSUM matrix for the original 46 rotamers.

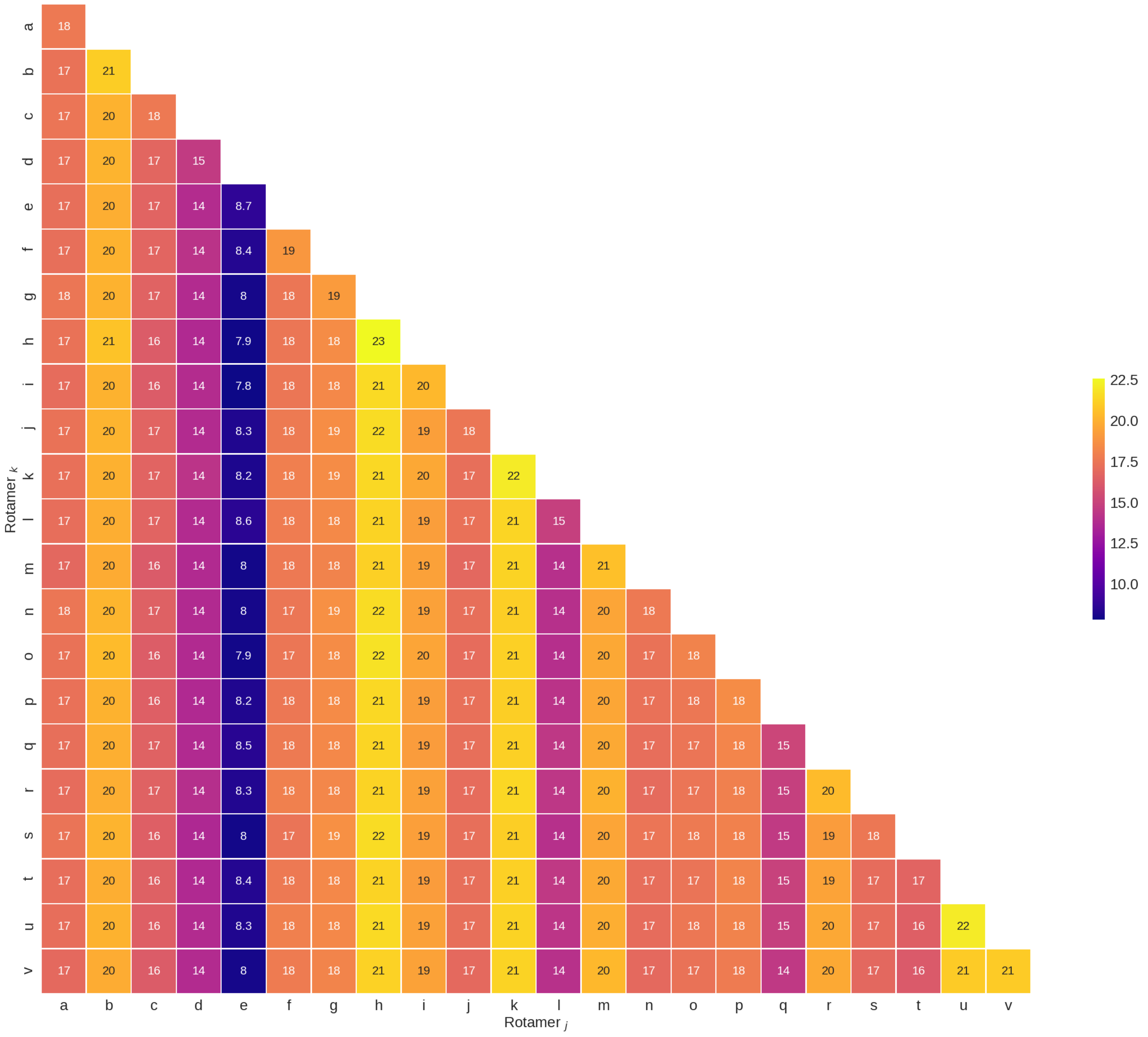


Figure S3: Representation of the ROSUM matrix for $\delta_{i-1}\delta_i\alpha\gamma$ families.

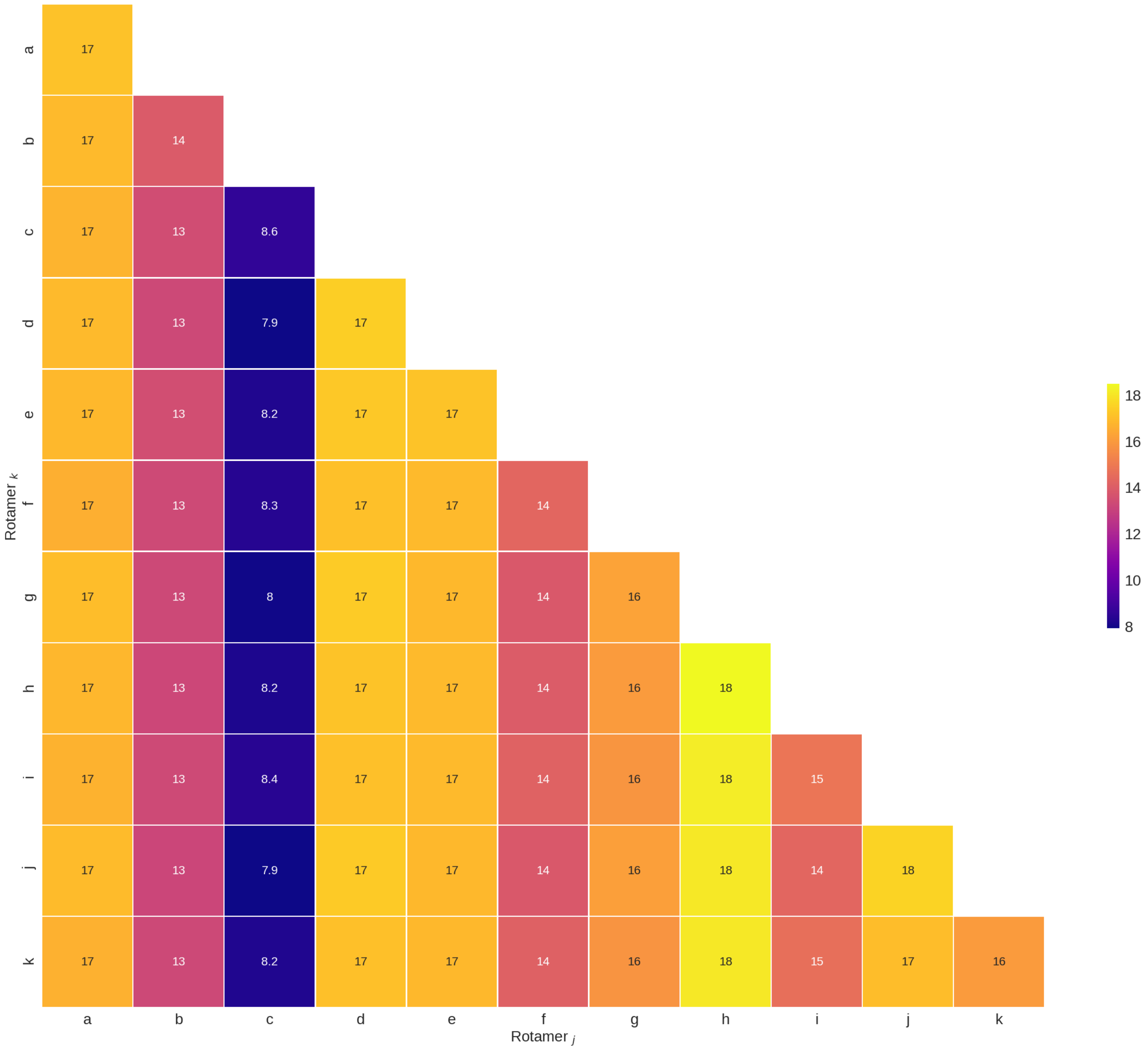


Figure S4: Representation of the ROSUM matrix for $\delta_{i-1}\delta_i\alpha$ families.

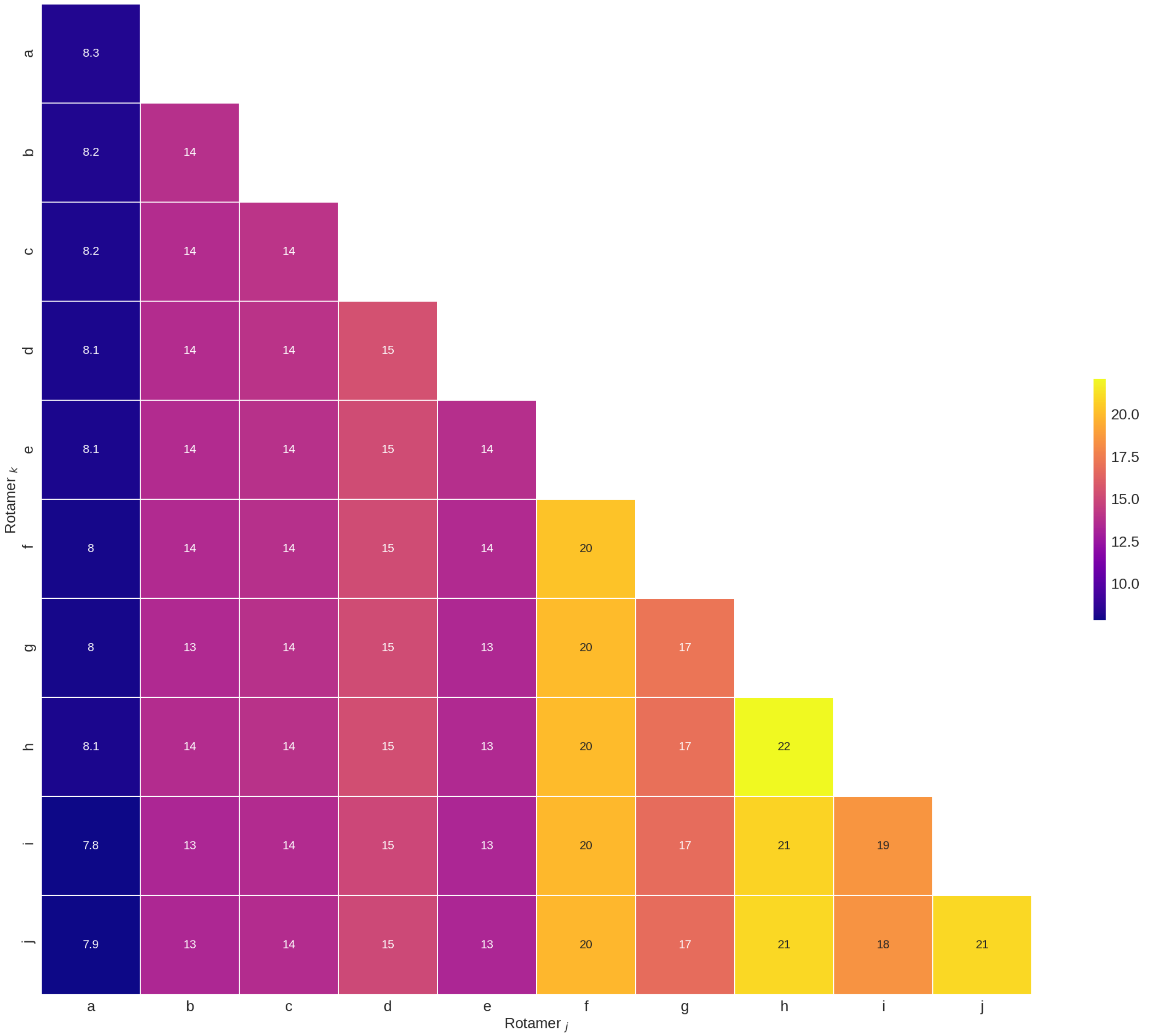


Figure S5: Representation of the ROSUM matrix for $\delta_{i-1}\delta_i\gamma$ families.

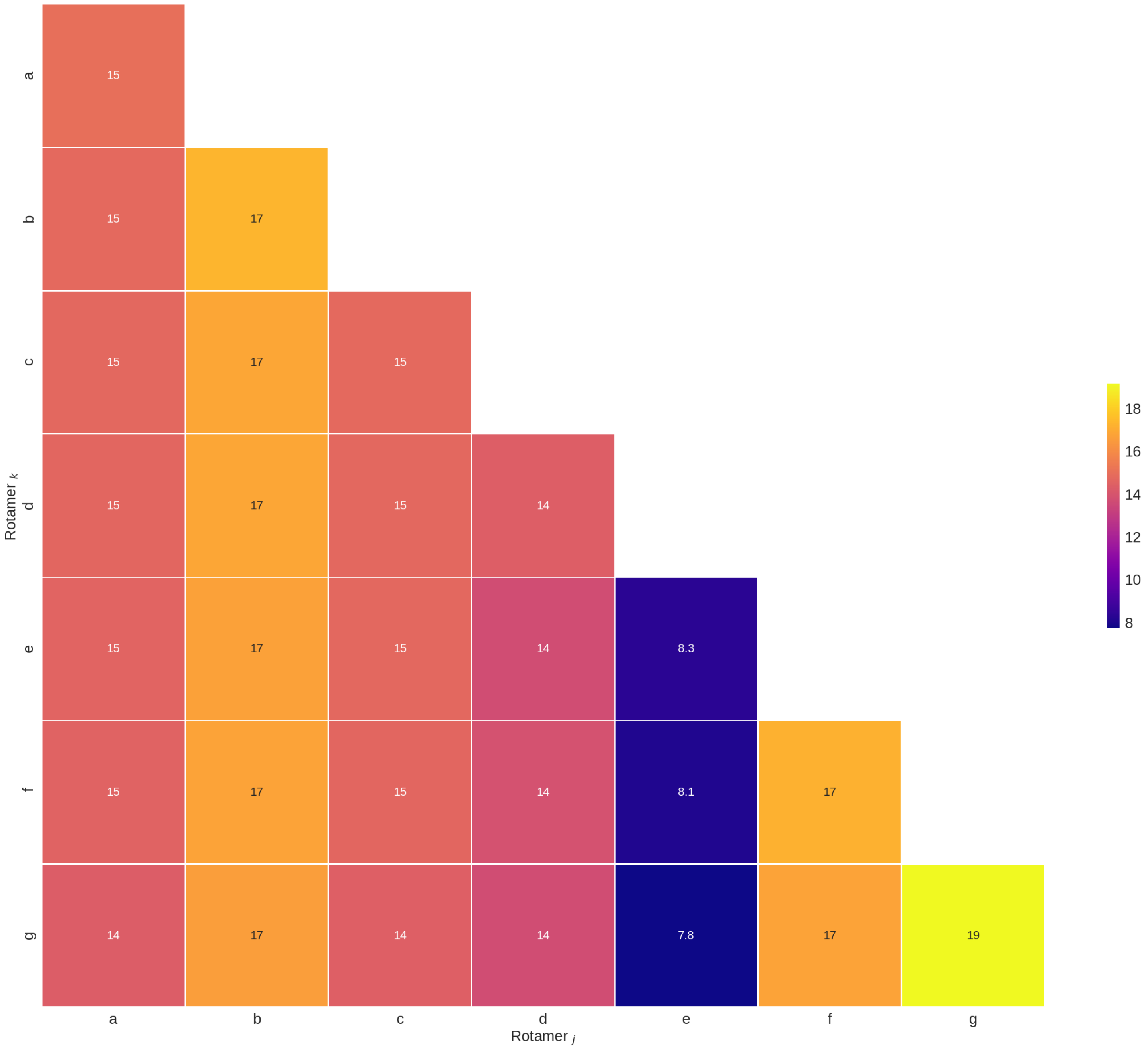


Figure S6: Representation of the ROSUM matrix for $\alpha\gamma$ families.

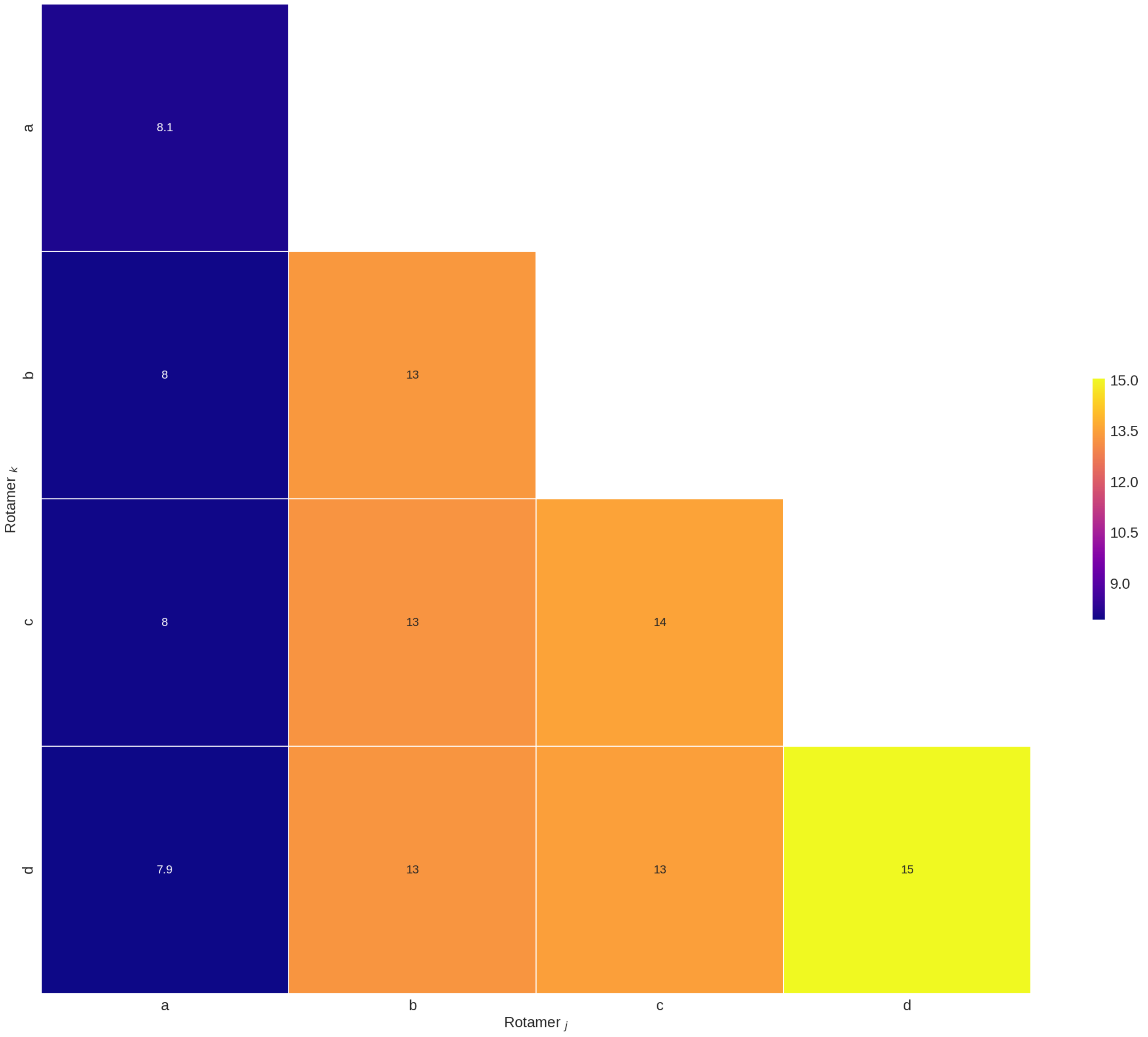


Figure S7: Representation of the ROSUM matrix for $\delta_{i-1}\delta_i$ families.

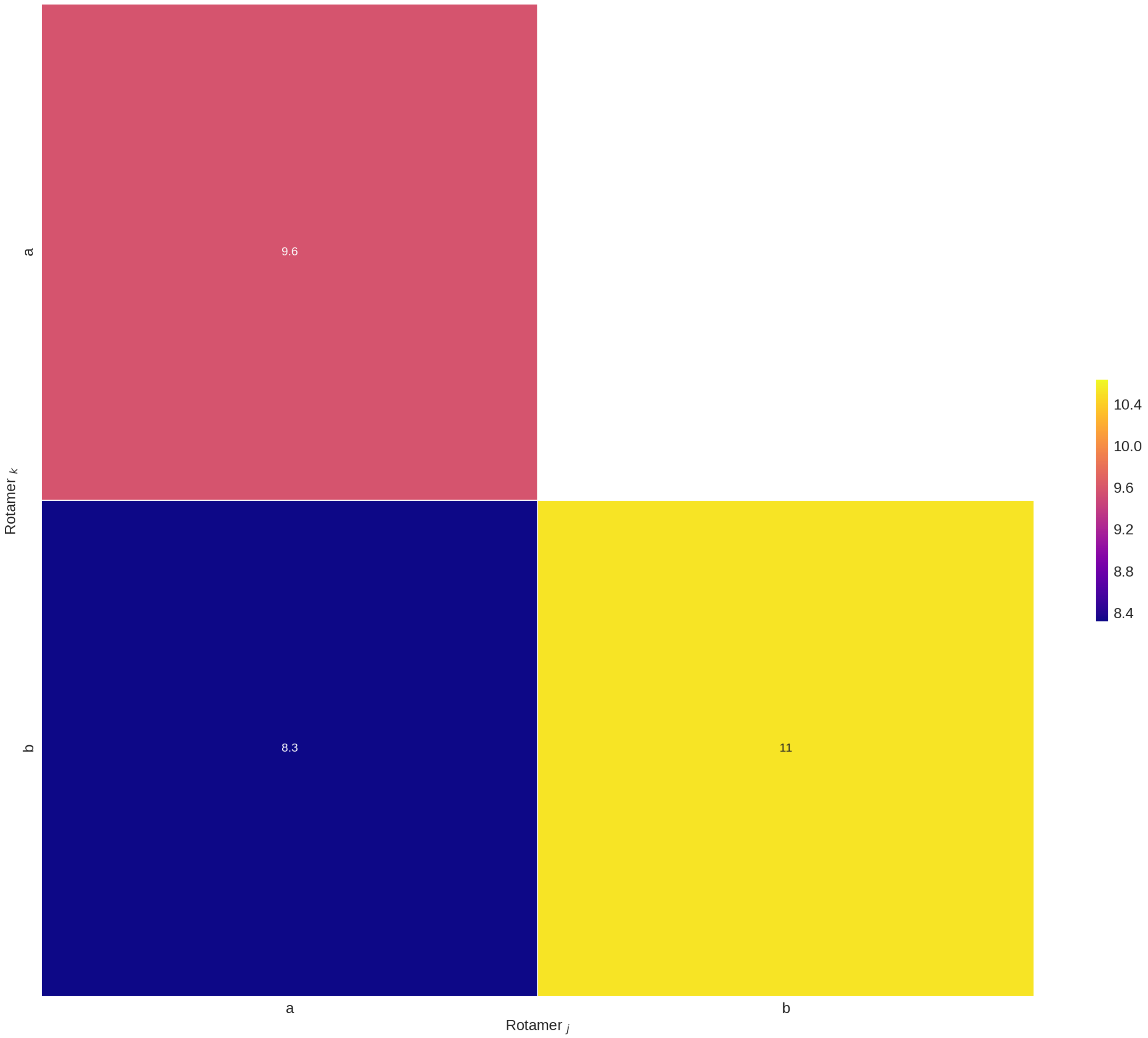


Figure S8: Representation of the ROSUM matrix for A_noA families.

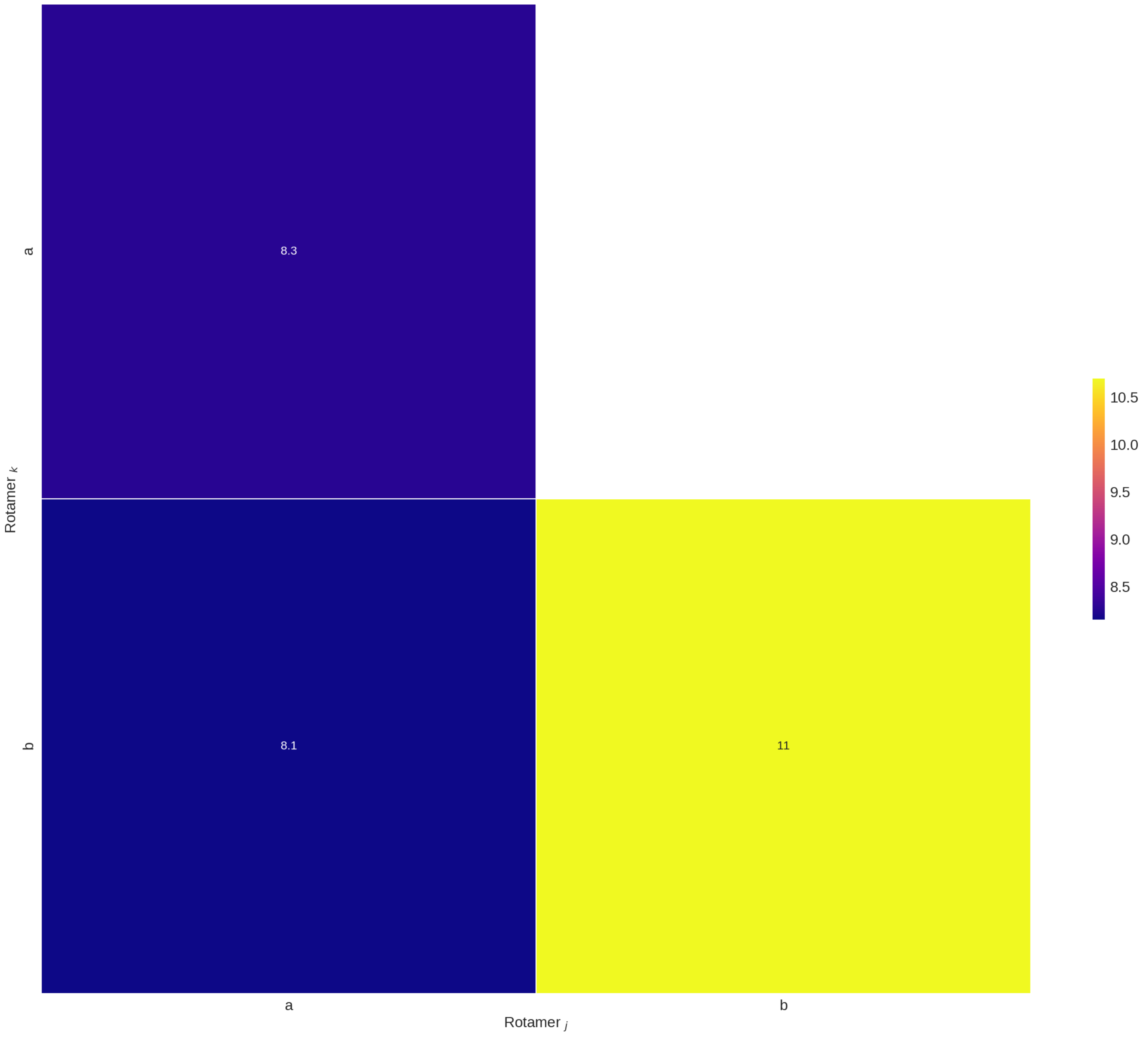


Figure S9: Representation of the ROSUM matrix for A*_noA* families.

### 4 Effective references

| Reference type | Value |
|---|---|
| **Simple** | 185.0 |
| **Sequence** | |
| AA | 181.5 |
| AC | 183.5 |
| AG | 185.0 |
| AU | 185.0 |
| CA | 185.0 |
| CC | 180.0 |
| CG | 187.9 |
| CU | 180.0 |
| GA | 185.0 |
| GC | 185.8 |
| GG | 185.2 |
| GU | 184.2 |
| UA | 180.0 |
| UC | 185.0 |
| UG | 189.7 |
| UU | 185.0 |
| **Puckering** | |
| C3'endo-C3'endo | 183.9 |
| C3'endo-C2'endo | 181.2 |
| C2'endo-C3'endo | 182.9 |
| C2'endo-C2'endo | 181.6 |
| **C_mean** | |
| C1'i-1 | 183.04 |
| C2'i-1 | 179.30 |
| C3'i-1 | 179.91 |
| C4'i-1 | 181.76 |
| C5'i-1 | 183.06 |
| C1'i | 182.67 |
| C2'i | 179.23 |
| C3'i | 180.30 |
| C4'i | 182.06 |
| C5'i | 183.36 |
| **C_opt** | |
| C1' | a= 74.75, b= 0.19 |
| C2' | a= 70.58, b= 0.06 |
| C3' | a= 56.64, b= 0.22 |
| C4' | a= 63.79, b= 0.23 |
| C5' | a= 53.75, b= 0.18 |

Table S2: Effective references used to transform the theoretical $^{13}$C shieldings into chemical shifts. The **sequence** and **puckering** correspond to the reference values that gave the highest performance in experimental vs. theoretical classification using the 1-NN model. The **C_mean** values were obtained by comparing the mean value of the distribution of experimental $^{13}$C chemical shifts and the mean value of the distribution of theoretical $^{13}$C shieldings, for the ten different nuclei in a dinucleotide (or suite). For **C_opt**, the values of a and b in the equation $\delta_{comp}$ = a + b*$\sigma_{comp}$ for the five different carbon nuclei in a nucleotide are given. Here, $\delta_{comp}$ and $\sigma_{comp}$ refer to theoretical chemical shifts and theoretical shieldings, respectively. The values of a and b were obtained trough a hierarchical model of a linear regression using PyMC3 [2] for a set of dinucleotides with theoretical and experimental $^{13}$C chemical shifts.

### 5 Classification models

#### 5.1 Scikit-learn parameters

| Model | Parameters |
|---|---|
| RandomForestClassifier | (criterion='gini', max_depth=20, n_estimators=10, max_features=1) |
| RandomForestClassifier | (criterion='gini', max_depth=20, n_estimators=10, max_features='auto') |
| RandomForestClassifier | (criterion='entropy', max_depth=20, n_estimators=10, max_features=1) |
| RandomForestClassifier | (criterion='entropy', max_depth=20, n_estimators=10, max_features='auto') |
| DecisionTreeClassifier | (criterion='gini', max_depth=20) |
| DecisionTreeClassifier | (criterion='gini', max_depth=None) |
| DecisionTreeClassifier | (criterion='entropy', max_depth=20) |
| DecisionTreeClassifier | (criterion='entropy', max_depth=None) |
| SVC | (kernel='linear', C=0.025) |
| SVC | (kernel='linear', C=0.1) |
| SVC | (kernel='linear', C=0.5) |
| SVC | (kernel='linear', C=1.0) |
| SVC | (kernel='linear', C=2.0) |
| SVC | (kernel='rbf', C=0.025) |
| SVC | (kernel='rbf', C=0.1) |
| SVC | (kernel='rbf', C=0.5) |
| SVC | (kernel='rbf', C=1.0) |
| SVC | (kernel='rbf', C=2.0) |
| KNeighborsClassifier | (k=1) |
| KNeighborsClassifier | (k=2) |
| KNeighborsClassifier | (k=3) |
| KNeighborsClassifier | (k=4) |
| KNeighborsClassifier | (k=5) |
| MLPClassifier | (alpha=0.0001, max_iter=500, solver='lbfgs') |
| MLPClassifier | (alpha=0.0001, max_iter=500, solver='sgd') |
| MLPClassifier | (alpha=0.0001, max_iter=500, solver='adam') |
| MLPClassifier | (alpha=0.0001, max_iter=500, solver='sgd') |
| MLPClassifier | (alpha=0.0001, max_iter=750, solver='sgd') |
| MLPClassifier | (alpha=0.0001, max_iter=1000, solver='sgd') |

Table S3: Sci-kit learn models and parameters used for classification.

#### 5.2 Scikit-learn parameters for the highest scores of experimental vs. theoretical classification

Table S4: Scikit–learn classifiers parameters for the highest values of weighted accuracy.

| **Rotamer families** | **Reference**[a] | **Classifier**[a] | **Scikit-learn parameters** | **Weighted accuracy** |
|---|---|---|---|---|
| 46 | Sequence | NN | KNeighborsClassifier(2) | 0.22 |
| $\delta_{i-1}\delta_i\alpha\gamma$ | Sequence | NN | KNeighborsClassifier(4) | 0.41 |
| $\delta_{i-1}\delta_i\alpha$ | Simple | NN | KNeighborsClassifier(4) | 0.46 |
| $\delta_{i-1}\delta_i\gamma$ | Simple | NN | KNeighborsClassifier(2) | 0.57 |
| $\alpha\gamma$ | Sequence | NN | KNeighborsClassifier(4) | 0.48 |
| $\delta_{i-1}\delta_i$ | Sequence | NN | KNeighborsClassifier(2) | 0.73 |
| A*_noA* | Sequence | NN | KNeighborsClassifier(1) | 0.64 |
| A _noA | Puckering | NN | KNeighborsClassifier(2) | 0.64 |
| 46 | Sequence | DT | DecisionTreeClassifier(criterion='entropy', max_depth=20) | 0.09 |
| $\delta_{i-1}\delta_i\alpha\gamma$ | Sequence | DT | DecisionTreeClassifier(criterion='entropy', max_depth=20) | 0.29 |
| $\delta_{i-1}\delta_i\alpha$ | Simple | DT | DecisionTreeClassifier(criterion='gini', max_depth=None) | 0.36 |
| $\delta_{i-1}\delta_i\gamma$ | Simple | DT | DecisionTreeClassifier(criterion='entropy', max_depth=None) | 0.56 |
| $\alpha\gamma$ | Sequence | DT | DecisionTreeClassifier(criterion='entropy', max_depth=None) | 0.35 |
| $\delta_{i-1}\delta_i$ | Simple | DT | DecisionTreeClassifier(criterion='entropy', max_depth=None) | 0.69 |
| A*_noA* | Sequence | DT | DecisionTreeClassifier(criterion='entropy', max_depth=20) | 0.61 |
| A _noA | Sequence | DT | DecisionTreeClassifier(criterion='entropy', max_depth=20) | 0.58 |
| 46 | Simple | RF | RandomForestClassifier(criterion='gini', max_depth=20, n_estimators=10, max_features=1) | 0.14 |
| $\delta_{i-1}\delta_i\alpha\gamma$ | Simple | RF | RandomForestClassifier(criterion='entropy', max_depth=20, n_estimators=10, max_features=1) | 0.36 |
| $\delta_{i-1}\delta_i\alpha$ | Simple | RF | RandomForestClassifier(criterion='entropy', max_depth=20, n_estimators=10, max_features=1) | 0.37 |
| $\delta_{i-1}\delta_i\gamma$ | Simple | RF | RandomForestClassifier(criterion='entropy', max_depth=20, n_estimators=10, max_features=1) | 0.56 |
| $\alpha\gamma$ | Sequence | RF | RandomForestClassifier(criterion='entropy', max_depth=20, n_estimators=10, max_features=1) | 0.41 |
| $\delta_{i-1}\delta_i$ | Puckering | RF | RandomForestClassifier(criterion='gini', max_depth=20, n_estimators=10, max_features=1) | 0.70 |
| A*_noA* | Sequence | RF | RandomForestClassifier(criterion='entropy', max_depth=20, n_estimators=10, max_features='auto') | 0.62 |
| A _noA | Sequence | RF | RandomForestClassifier(criterion='entropy', max_depth=20, n_estimators=10, max_features='auto') | 0.56 |
| 46 | Puckering | MLP | MLPClassifier(alpha=0.0001, max_iter=500, solver='lbfgs') | 0.06 |
| $\delta_{i-1}\delta_i\alpha\gamma$ | C_opt | MLP | MLPClassifier(alpha=0.0001, max_iter=750, solver='sgd') | 0.51 |
| $\delta_{i-1}\delta_i\alpha$ | C_opt | MLP | MLPClassifier(alpha=0.0001, max_iter=1000, solver='sgd') | 0.45 |
| $\delta_{i-1}\delta_i\gamma$ | Simple | MLP | MLPClassifier(alpha=0.0001, max_iter=1000, solver='sgd') | 0.55 |
| $\alpha\gamma$ | Puckering | MLP | MLPClassifier(alpha=0.0001, max_iter=1000, solver='sgd') | 0.60 |
| $\delta_{i-1}\delta_i$ | Simple | MLP | MLPClassifier(alpha=0.0001, max_iter=500, solver='adam') | 0.65 |
| A*_noA* | C_opt | MLP | MLPClassifier(alpha=0.0001, max_iter=500, solver='lbfgs') | 0.57 |
| A _noA | C_opt | MLP | MLPClassifier(alpha=0.0001, max_iter=500, solver='lbfgs') | 0.52 |
| 46 | Sequence | SVM | SVC(kernel='linear', C=0.025) | 0.20 |
| $\delta_{i-1}\delta_i\alpha\gamma$ | Simple | SVM | SVC(kernel='rbf', C=0.025) | 0.51 |
| $\delta_{i-1}\delta_i\alpha$ | Simple | SVM | SVC(kernel='rbf', C=0.025) | 0.47 |
| $\delta_{i-1}\delta_i\gamma$ | Simple | SVM | SVC(kernel='rbf', C=0.025) | 0.55 |
| $\alpha\gamma$ | Simple | SVM | SVC(kernel='rbf', C=0.025) | 0.60 |
| $\delta_{i-1}\delta_i$ | Simple | SVM | SVC(kernel='linear', C=0.025) | 0.71 |
| A*_noA* | Simple | SVM | SVC(kernel='rbf', C=1.0) | 0.53 |
| A _noA | Simple | SVM | SVC(kernel='linear', C=0.5) | 0.62 |

***a*** **Reference** refers to the effective references used to obtain the theoretical CS (see Main text).
***b*** **Classifier**: Nearest Neighbor (NN), Decision Tree (DT), Random Forest (RF), Multi-Layer Perceptron (MLP) and Support Vector Machine (SVM).

Table S5: Scikit–learn classifiers parameters for the highest values of $F_1$ score.

| **Rotamer families** | **Reference**[a] | **Classifier**[b] | **Scikit-learn parameters** | **F1 score** |
|---|---|---|---|---|
| 46 | Sequence | NN | KNeighborsClassifier(2) | 0.34 |
| $\delta_{i-1}\delta_i\alpha\gamma$ | Simple | NN | KNeighborsClassifier(1) | 0.53 |
| $\delta_{i-1}\delta_i\alpha$ | Simple | NN | KNeighborsClassifier(1) | 0.55 |
| $\delta_{i-1}\delta_i\gamma$ | Simple | NN | KNeighborsClassifier(1) | 0.63 |
| $\alpha\gamma$ | Sequence | NN | KNeighborsClassifier(4) | 0.59 |
| $\delta_{i-1}\delta_i$ | Simple | NN | KNeighborsClassifier(1) | 0.77 |
| A*_noA* | Sequence | NN | KNeighborsClassifier(1) | 0.63 |
| A _noA | Sequence | NN | KNeighborsClassifier(2) | 0.61 |
| 46 | Simple | DT | DecisionTreeClassifier(criterion='entropy', max_depth=None) | 0.16 |
| $\delta_{i-1}\delta_i\alpha\gamma$ | Simple | DT | DecisionTreeClassifier(criterion='gini', max_depth=20) | 0.43 |
| $\delta_{i-1}\delta_i\alpha$ | Simple | DT | DecisionTreeClassifier(criterion='gini', max_depth=None) | 0.48 |
| $\delta_{i-1}\delta_i\gamma$ | Simple | DT | DecisionTreeClassifier(criterion='gini', max_depth=None) | 0.61 |
| $\alpha\gamma$ | Sequence | DT | DecisionTreeClassifier(criterion='entropy', max_depth=20) | 0.47 |
| $\delta_{i-1}\delta_i$ | Simple | DT | DecisionTreeClassifier(criterion='gini', max_depth=None) | 0.73 |
| A*_noA* | Sequence | DT | DecisionTreeClassifier(criterion='entropy', max_depth=20) | 0.59 |
| A _noA | Sequence | DT | DecisionTreeClassifier(criterion='entropy', max_depth=20) | 0.52 |
| 46 | Simple | RF | RandomForestClassifier(criterion='gini', max_depth=20, n_estimators=10, max_features=1) | 0.24 |
| $\delta_{i-1}\delta_i\alpha\gamma$ | Simple | RF | RandomForestClassifier(criterion='entropy', max_depth=20, n_estimators=10, max_features=1) | 0.49 |
| $\delta_{i-1}\delta_i\alpha$ | Puckering | RF | RandomForestClassifier(criterion='entropy', max_depth=20, n_estimators=10, max_features=1) | 0.49 |
| $\delta_{i-1}\delta_i\gamma$ | Puckering | RF | RandomForestClassifier(criterion='entropy', max_depth=20, n_estimators=10, max_features='auto') | 0.64 |
| $\alpha\gamma$ | C_mean | RF | RandomForestClassifier(criterion='entropy', max_depth=20, n_estimators=10, max_features=1) | 0.53 |
| $\delta_{i-1}\delta_i$ | Puckering | RF | RandomForestClassifier(criterion='gini', max_depth=20, n_estimators=10, max_features=1) | 0.74 |
| A*_noA* | Sequence | RF | RandomForestClassifier(criterion='entropy', max_depth=20, n_estimators=10, max_features='auto') | 0.61 |
| A _noA | Sequence | RF | RandomForestClassifier(criterion='entropy', max_depth=20, n_estimators=10, max_features='auto') | 0.49 |
| 46 | Puckering | MLP | MLPClassifier(alpha=0.0001, max_iter=500, solver='lbfgs') | 0.11 |
| $\delta_{i-1}\delta_i\alpha\gamma$ | Sequence | MLP | MLPClassifier(alpha=0.0001, max_iter=750, solver='sgd') | 0.55 |
| $\delta_{i-1}\delta_i\alpha$ | Puckering | MLP | MLPClassifier(alpha=0.0001, max_iter=1000, solver='sgd') | 0.52 |
| $\delta_{i-1}\delta_i\gamma$ | C_mean | MLP | MLPClassifier(alpha=0.0001, max_iter=500, solver='lbfgs') | 0.62 |
| $\alpha\gamma$ | Sequence | MLP | MLPClassifier(alpha=0.0001, max_iter=1000, solver='sgd') | 0.63 |
| $\delta_{i-1}\delta_i$ | Simple | MLP | MLPClassifier(alpha=0.0001, max_iter=500, solver='adam') | 0.72 |
| A*_noA* | C_opt | MLP | MLPClassifier(alpha=0.0001, max_iter=500, solver='lbfgs') | 0.44 |
| A _noA | C_opt | MLP | MLPClassifier(alpha=0.0001, max_iter=500, solver='lbfgs') | 0.41 |
| 46 | Sequence | SVM | SVC(kernel='linear', C=0.025) | 0.31 |
| $\delta_{i-1}\delta_i\alpha\gamma$ | Simple | SVM | SVC(kernel='rbf', C=0.025) | 0.55 |
| $\delta_{i-1}\delta_i\alpha$ | C_mean | SVM | SVC(kernel='rbf', C=0.025) | 0.57 |
| $\delta_{i-1}\delta_i\gamma$ | C_mean | SVM | SVC(kernel='linear', C=0.025) | 0.6 |
| $\alpha\gamma$ | Simple | SVM | SVC(kernel='rbf', C=0.025) | 0.62 |
| $\delta_{i-1}\delta_i$ | Simple | SVM | SVC(kernel='linear', C=0.025) | 0.75 |
| A*_noA* | Simple | SVM | SVC(kernel='rbf', C=2.0) | 0.36 |
| A _noA | Simple | SVM | SVC(kernel='linear', C=0.5) | 0.62 |

***a*** **Reference** refers to the effective references used to obtain the theoretical CS (see Main text).

***b*** **Classifier**: Nearest Neighbor (NN), Decision Tree (DT), Random Forest (RF), Multi-Layer Perceptron (MLP) and Support Vector Machine (SVM)
